# Reducing peptide sequence bias in quantitative mass spectrometry data with machine learning

**DOI:** 10.1101/2022.04.11.487945

**Authors:** Ayse Dincer, Yang Lu, Devin Schweppe, Sewoong Oh, William Stafford Noble

## Abstract

Quantitative mass spectrometry measurements of peptides necessarily incorporate sequence-specific biases that reflect the behavior of the peptide during enzymatic digestion, liquid chromatography, and in the mass spectrometer. These sequence-specific effects impair quantification accuracy, yielding peptide quantities that are systematically under- or over-estimated. We provide empirical evidence for the existence of such biases, and we use a deep neural network, called Pepper, to automatically identify and reduce these biases. The model generalizes to new proteins and new runs within a related set of MS/MS experiments, and the learned coefficients themselves reflect expected physicochemical properties of the corresponding peptide sequences. The resulting adjusted abundance measurements are more correlated with mRNA-based gene expression measurements than the unadjusted measurements. Pepper is suitable for data generated on a variety of mass spectrometry instruments, and can be used with labeled or label-free approaches, and with data-independent or data-dependent acquisition.

## 1 Introduction

Tandem mass spectrometry (MS/MS) can be used to quantify thousands of peptides in a complex biological mixture. Regardless of how the quantitative values are extracted from the data—using labeling strategies such as iTRAQ or TMT, peak areas from precursor scans, or spectral counts—all such quantitative measurements exhibit biases. In general, “bias” in our context means that quantitative measurements from a mass spectrometry experiment are systematically skewed, either positively or negatively, relative to the true abundance of the measured molecular species. Some of these biases depend in part on properties of the peptide sequence, such as how susceptible the peptide is to enzymatic cleavage, how efficiently the peptide traverses the liquid chromatography column, how easily the peptide ionizes in electrospray, and how easily and uniformly the peptide fragments in the dissociation phase of the MS/MS.

Numerous efforts have been made to identify and quantify these sequence-specific effects [1–7]. For example, a protein’s “proteotypic peptides”—peptides that are repeatedly and consistently identified for a given protein—can be identified using machine learning methods that take into account a wide range of physicochemical properties of amino acids, including charge, secondary structure propensity and hydrophobicity. Peptide hydrophobicity, in particular, strongly affects ionization in electrospray settings [8, 9].

In this work, we focus on biases that arise directly from the amino acid sequence itself. Many biases exist that our approach is not designed to address. This includes, for example, the effect of secondary or tertiary protein structure, which could inhibit cleavage by trypsin, as well as competitive effects due to chromatographic coelution of other peptides. We focus on sequence-induced biases because our machine learning framework depends on peptide-level labels, as described below.

Our goal is to train a machine learning model to quantify these peptide-specific properties, with the aim of adjusting the observed quantities to remove these effects. Our approach rests on two primary assumptions. First, we assume that each peptide is measured in its linear dynamic range and that the observed measurement *q_ik_* of peptide *i* in run *k* can be decomposed into a peptide coefficient *c_i_* and an adjusted peptide abundance *α_ik_* such that *q_ik_* = *c_i_α_ik_*. Second, we assume that unique sibling peptides, i.e., peptides that occur exactly once in the protein database and that co-occur on a given protein sequence, should have equal abundances within a single MS/MS run. We use these assumptions to train a deep neural network, Pepper, that takes as input a peptide sequence *p_i_* and charge state *z_i_* and produces as output the corresponding peptide coefficient *c_i_*, thereby revealing the adjusted peptide abundance *α_ik_*.

We demonstrate that, by removing peptide-specific effects from the observed MS/MS quantities, the adjusted abundance values *α_ik_* provide more accurate abundance measurements than the observed values. First, we provide empirical evidence for the consistency of peptide coefficients inferred from disjoint sets of runs, and we show that a Pepper model trained to predict these coefficients can generalize to new peptides in new runs. We demonstrate the robustness of the approach to the type of noise that one would expect to arise from the presence of proteoforms in the sample that are not represented in the reference proteome database. We show that the learned coefficients exhibit significant correlation with several key peptide properties, in agreement with previous research [1–7]. We also show that the adjusted peptide abundances yield a small but highly significant improvement in the degree of correlation between MS/MS and mRNA-based gene expression measurements. We provide open source code for training models from MS/MS data, which can be used to de-bias any given matrix of peptide-level abundances (http://github.com/Noble-Lab/Pepper).

### 1.1 Related work

The measurements produced by a mass spectrometry experiment invariably exhibit biases. In particular, among the many thousands of distinct tryptic peptides in a complex biological mixture, only a small fraction are typically observed [10], and some peptides are preferentially identified regardless of whether they occur on the most abundant proteins [1]. These commonly observed peptides are called *proteotypic peptides*, and automatically identifying them can be important for accurate protein detection and quantification [1]. In particular, better understanding of the biases underlying proteotypicity can lead to changes in experimental design, and knowing the potentially observable peptides for a protein beforehand might increase our confidence in the identification of missing proteins in a sample [2]. Accordingly, numerous methods have been developed to predict proteotypic peptides using machine learning.

The first such method used physicochemical properties of amino acids summarized at the peptide level and selected properties that can most successfully differentiate observed versus unobserved peptides using Kolmogorov-Smirnov and Kullback-Leibler distances [1]. Using these most discriminative features, Mallick *et al*. fitted a Gaussian mixture likelihood function to predict the probability of detection for each peptide, thereby achieving a test accuracy above 85%. The trained model showed robust performance across various datasets and organisms.

Thereafter, a series of machine learning methods were developed to improve the accuracy of proteotypic peptide prediction and to target the predictions to particular applications (Table 1). Sanders *et al*. [2] developed a neural network-based predictor, which can encode non-linear relations among the input features. Similarly, Webb-Robertson et *al*. [3] developed a non-linear support vector machine classifier to predict proteotypic peptides, after first applying the Fisher Criterion Score to select a subset of peptide features to train on. Unlike previous approaches, they jointly predict proteotypicity (i.e., detection probability) and the elution time of a peptide. Several methods focused on predicting which peptides will be most useful in a targeted MS setting. In one such study, Fusaro *et al*. used a random forest classifier for the prediction of high-responding peptides to be used as targets [4], and Searle et al.’s PREGO model [6] adopted a similar approach, using data independent acquisition (DIA) fragment intensities to define their training set. The Peptide Prediction with Abundance (PPA) method also uses a neural network and predicts not only the probability of detection for the peptide but also the corresponding protein abundance that would enable detecting that peptide [7]. Finally, CONSeQuence uses a consensus machine learning approach—an ensemble of support vector machine, random forest, genetic programming, and neural network models—to select peptides for absolute quantification experiments [5].

**Table 1:** Methods for predicting proteotypic peptides.

|  | Year | Model | Experiment type |
| --- | --- | --- | --- |
| Mallick *et al.* [1] | 2006 | Gaussian mixture model | DDA |
| Sanders *et al.* [2] | 2007 | Neural network | DDA |
| Webb-Robertson *et al.* [3] | 2008 | Support vector machine | DDA |
| Fusaro *et al.* [4] | 2009 | Random forest | Targeted MS |
| CONSeQuence [5] | 2011 | Ensemble model | Absolute quantification |
| PREGO [6] | 2015 | Neural network | Targeted MS |
| PPA [7] | 2015 | Neural network | DDA |

All of the studies cited above have focused on identifying proteotypic or high-responding peptides. In contrast, our goal is to quantify the peptide biases in quantitative MS experiments. Thus, rather than classifying the peptides as low-responding versus high-responding, we aim to learn peptide coefficients that can allow us to reduce bias in our measurements. To our knowledge, ours is the first machine learning approach to quantifying sequence-induced bias for mass spectrometry.

Another key point that differentiates our work from the previous studies is that instead of using the physicochemical properties of the amino acids, we take the peptide sequence itself as input. Using the physicochemical properties of peptides has many intrinsic limitations. Most importantly, all previous studies used the amino acid properties summarized at the peptide level (e.g., using the mean or sum) which results in losing sequence-specific information. In practice, small changes in the order of amino acids within a peptide sequence might significantly affect how the peptide behaves in a mass spectrometer. Therefore, our model can potentially explain biases that a model trained solely on summary-level peptide properties might fail to capture. Our sequence-based approach has the added advantage that we do not need to worry about pre-processing large tables of peptide features in order to reduce redundancy, as was done in many previous studies [1–3, 5, 6].

## 2 Methods

### 2.1 A neural network model for predicting peptide coefficients

We designed a neural network architecture, Pepper, that aims to learn peptide coefficients from quantitative mass spectrometry measurements (Figure 1). The inputs to Pepper are a matrix of measured peptide abundances and corresponding databases of peptides and proteins. Say that we are given a matrix *Q* of peptide measurements, where *q_ik_* is the observed abundance of peptide *i* in run *k*. We column normalize *Q* so that the sum of all abundances within a given run is constant across runs. We are also given a peptide database *P* and corresponding protein database 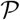. For the purposes of training our model, we preprocess *P* to eliminate all peptides that appear in more than one protein. In addition, to reduce issues related to unexpected proteoforms, we identify any peptide that contains a variable modification, such as phosphorylation or oxidation of methionine, and we eliminate both the modified and unmodified form of the peptide from *P*. Similarly, we eliminate all peptides that overlap one another, due to missed or non-enzymatic cleavages. Finally, we retain in 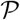 only proteins that contain at least two unique peptides.

**Figure 1:**
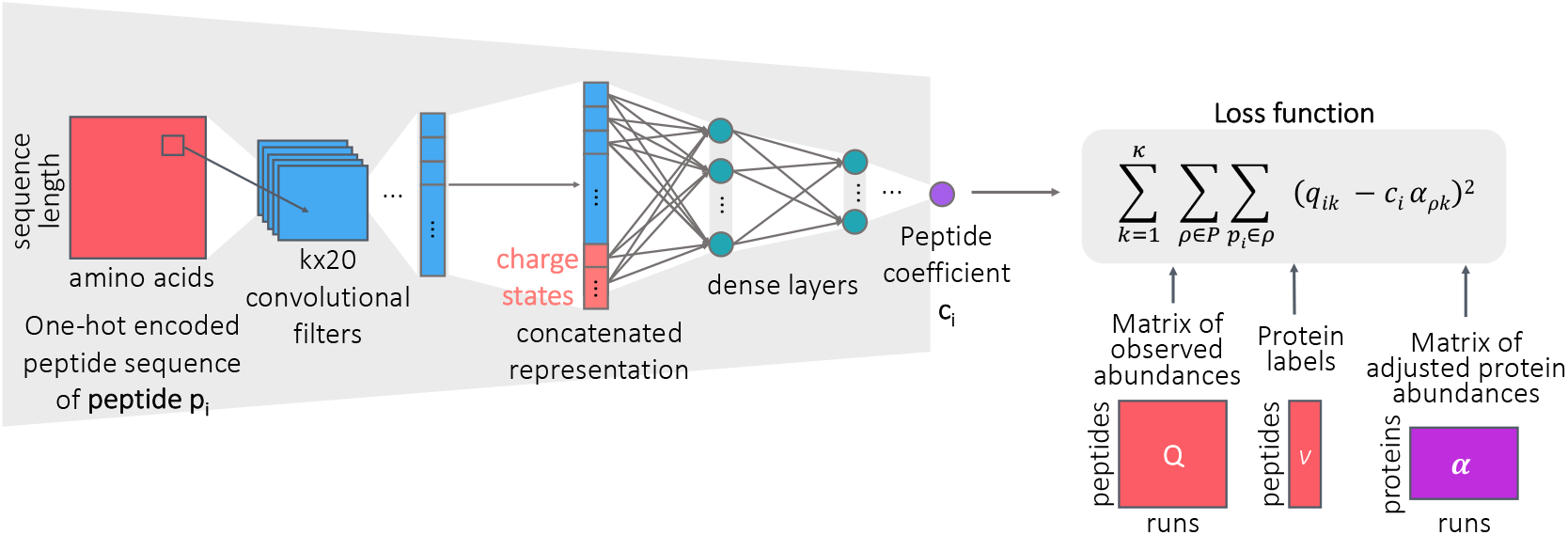
Peptide coefficient predictor. The neural network architecture for predicting peptide coefficients. The network takes as input the one-hot encoded peptide sequence and the charge state, and runs through convolutional layers followed by densely connected layers to output a peptide coefficient.

Pepper takes as input one-hot encoded peptide sequence, *p_i_*, along with the charge state, and produces as output the corresponding coefficient *c_i_* (i.e., *f*(*p_i_*)). The network is then trained using a loss function that captures our assumption that the adjusted abundances of all sibling peptides should be equal to one another and, thus, should be equal to the adjusted abundance of the corresponding protein, *ρ*:

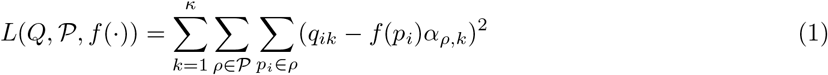

where *α_ρ,k_* is the adjusted abundance of protein *ρ* in run *k* and *κ* is the number of runs in *Q*. The model is trained subject to the constraints ∀*i c_i_* > 0 and ∀*_ρ,k_ α_ρ,k_* > 0. Note that the resulting coefficients can be used to adjust the measured abundances of all peptides, not just unique peptides, via 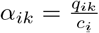.

We note that the elements of the *α* matrix are parameters of our model which we optimize along with the network weights while training the model. When calculating the loss function in Equation 1, we initialize the *α* matrix to the median observed peptide abundance per protein. To obtain the final adjusted protein abundance matrix 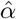 for a test set, we optimize over the fixed set of *Q* matrix and predicted *c_i_* values, and we use the resulting 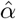 to calculate the final loss for the test set.

### 2.2 Data sets

The primary dataset was obtained from Guo *et al*. [11]. The data was generated from NCI-60 cancer cell lines using the PCT-SWATH workflow. Two replicates were obtained for each cell line, resulting in a total of 120 runs. SWATH-MS acquisition was used in a Sciex TripleTOF 5600 mass spectrometer with 32 windows of isolation width of 25. Peptides were detected at a peptide-level FDR threshold of 1% with OpenSWATH, using a human cancer cell line spectral library containing 86,209 proteotypic peptide precursors.

The other datasets were selected to reflect a range of instrument types, experimental protocols, and quantification methods (Table 2). The Slevlek *et al*. and Thomas *et al*. datasets provided peptide-level matrices of intensities. The CPTAC datasets were processed to obtain peptide quantities from the PSM files by repeating the preprocessing pipeline implemented by the CPTAC consortium, which starts with selecting the highest observed intensity for each sample and peptide across all fractions. Each sample was analyzed in 24 or 25 fractions, depending on the dataset. For each peptide, we selected the PSM with the highest “TotalAb” (i.e., total intensity across all TMT channels) among all fractions. The result is a peptide-by-sample matrix of measured intensities.

**Table 2:**
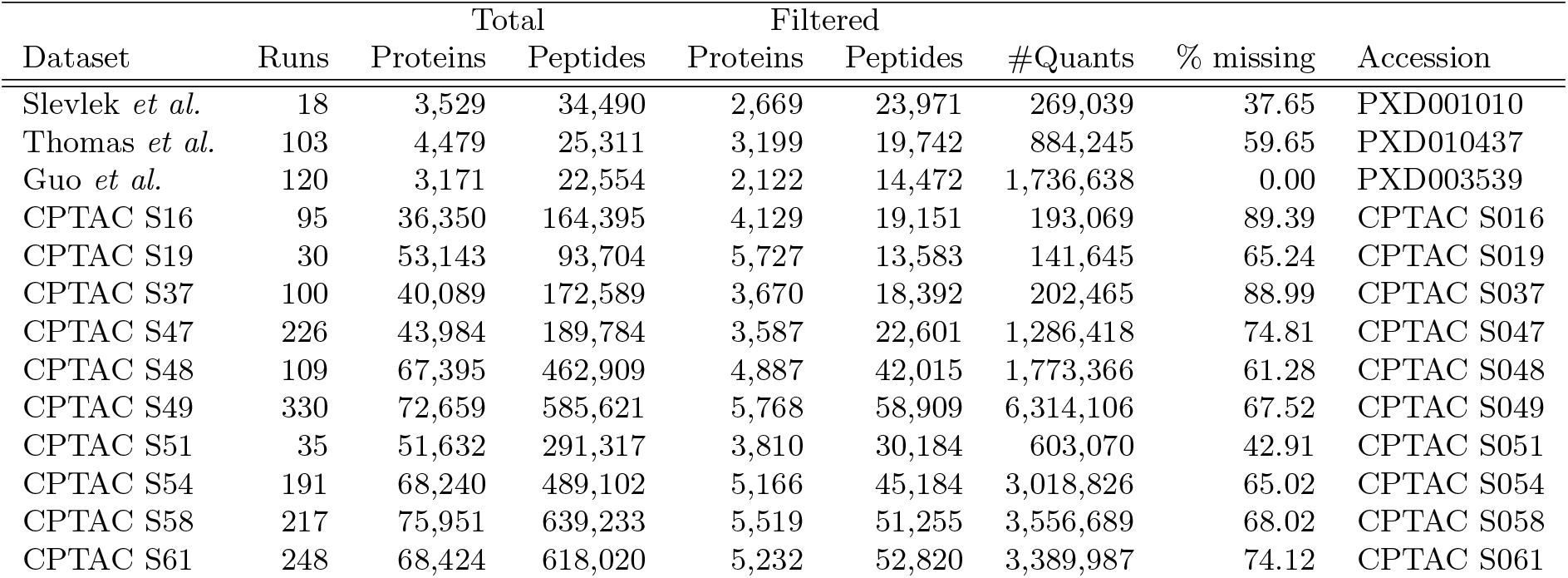
Datasets. The table lists datasets used in the study. The numbers of peptides and proteins are given before (“Total”) and after (“Filtered”) eliminating shared peptides and peptides with missed cleavages, peptides that occur in modified and unmodified forms, and peptides with no siblings.

For each data set, we normalize the peptide measurements so that the sum of peptide abundances is equal across all runs.

### 2.3 Filtering peptides

For a given quantitative matrix, we construct a filtered set of unique peptides by reducing the number of rows (peptides) in the matrix in four steps. First, we identify all peptides that occur in more than one protein, and we eliminate these from the matrix. Second, we identify all peptides that occur in both modified and unmodified forms, and we eliminate these from the matrix. Third, we eliminate all pairs of peptides that overlap one another due to missed cleavages. Fourth, among the remaining peptides, we identify and remove singletons, i.e., peptides with no siblings. The remaining peptides comprise the set of “filtered” peptides (Table 2). Note that if a peptide occurs in more than one charge state, these are treated as distinct peptides.

For input to the model, each peptide is encoded in a 1206-dimensional vector. The first six dimensions represent a one-hot encoding of the charge state (from +1 to +6). The remaining 1200 dimensions represent the peptide sequence, where each position is encoded with a 20-dimensional vector corresponding to different amino acids, up to a maximum length of 60.

### 2.4 Train and test set construction

To construct the train/test split, we split the data along two axes. First, we randomly segregate the runs in a ratio of 80%/20%. In this step, if a dataset contains replicate sets of runs, we make sure to keep the replicate runs within the same set. Second, we identify all proteins that contain at least one pair of sibling peptides, and we segregate the proteins into train and test sets in the same ratio.

The training set is comprised of all peptides that occur in training proteins, using measurements drawn from the training set runs. Conversely, the test set contains measurements of peptides within test proteins and measured in test runs. We also defined a validation set with the same process by splitting the training set into training and validation runs with a ratio of 90%/10%. We used this validation set for optimizing our deep learning model, such as hyperparameter tuning and early stopping.

### 2.5 Neural network architecture and training

The Pepper network takes as input the one-hot encoded peptide sequence and charge states and outputs a peptide coefficient. The neural network consists of a 2D convolutional layer containing 20 × *k* filters, each of which effectively extracts *k*-mers from the sequence. The output is flattened and concatenated with the charge state, which is then passed to the dense layers. ReLu activaton is used in each layer, and dropout layers are included after every layer. Pepper is trained using the Adam optimizer with an initial learning rate of 0.001 with gradient normalization. Early stopping on the validation set is used with a patience of 100 epochs and a threshold of 0.01 improvement in the loss. The best model, as measured by the loss function (Equation 1) on the validation set, is recovered when the training is done.

When calculating the loss function to update the model, we also take into account the protein labels *P* and the peptide-level measurements *Q*; however, these are not inputs to the model and thus are not used once the model has been trained. Note that missing values or zeros in the *Q* matrix are excluded from the loss calculation. The calculated loss values are normalized by dividing them by the total number of peptides and runs, which enables direct comparison between training, validation, and test sets.

Along with the dense and convolutional layers, we also define the *α* matrix as one of the parameters of our model, which we update while optimizing our loss function. We initialize the *α* matrix as the median abundance per protein.

For hyperparameter tuning, we used a random search over a grid of hyperparameters. Specifically, we sampled from a grid of filter size (3, 4, 5, 6, 7), number of filters (5, 10, 20, 40), number of layers (1, 2, 3, 4, 5), number of hidden nodes (10, 20, 40, 80), dropout rate (0.0, 0.25, 0.5, 0.75), learning rate (5e-4, 1e-3), and batch size (500, 1000), selecting the hyperparameters that yield the lowest loss (Equation 1) on the validation set. The selected hyperparameters were 10 filters each with a size of 3, a total of 4 hidden layers with 40 nodes in each dense layer, trained with a learning rate of 0.001, batch size of 1000, and a dropout rate of 0.25.

We repeated the hyperparameter selection procedure for each dataset separately, extending the grid when necessary, and selected the optimal hyperparameters. We used the same model architecture for all the CPTAC TMT11 datasets. We implemented our model using Keras with a Tensorflow backend.

## 3 Results

### 3.1 Empirical investigation of peptide coefficients

Prior to training a machine learning model to predict peptide coefficients from sequence, we investigated whether we observe consistency of quantities between pairs of sibling peptides across multiple MS runs. For this and most of the remaining experiments, we used SWATH-MS data obtained from the NCI-60 cancer cell lines [11]. The data consist of a total of 14,472 peptides and 120 runs, where two replicate runs were carried out for each of the 60 cancer cell lines. We made a list of all peptides identified in any one of the runs, and we eliminated shared peptides (i.e., peptides that appear in more than one protein) from the list. We then randomly selected a pair of unique peptides (*p_i_*, *p_j_*) and a random pair of runs (*r_k_*, *r_l_*). If both peptides were observed in both runs, then we recorded the corresponding observed quantities (*q_ik_*, *q_il_*, *q_jk_*, *q_jl_*). We assume that each of these quantities can be decomposed into a peptide coefficient and an adjusted abundance; e.g., *q_ik_* = *c_i_α_ik_*. Furthermore, we hypothesize that the adjusted abundances for a sibling peptide pair should be approximately equal to one another; i.e., if *p_i_* and *p_j_* are siblings, then *α_ik_* ≈ *α_jk_* and *α_il_* ≈ *α_jl_*. It follows, therefore, that the ratio of the abundances for the two sibling peptides between the two runs should be approximately equal:

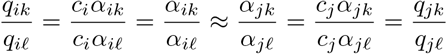

By contrast, we do not expect to observe any correspondence between ratios of pairs of non-sibling peptides.

To test our hypothesis, we repeated this sampling procedure many times and segregated the observed cross-run ratios into sibling and non-sibling pairs. The results of these analysis show evidence for consistency of peptide coefficients across runs (Figure 2): the Pearson correlation between intensity ratios of peptides are 0.39 and 0.03 for sibling and non-sibling peptides, respectively. This high correlation between ratios of sibling peptides demonstrates that sibling peptide measurements can be used as anchors for quantifying the peptide biases.

**Figure 2:**
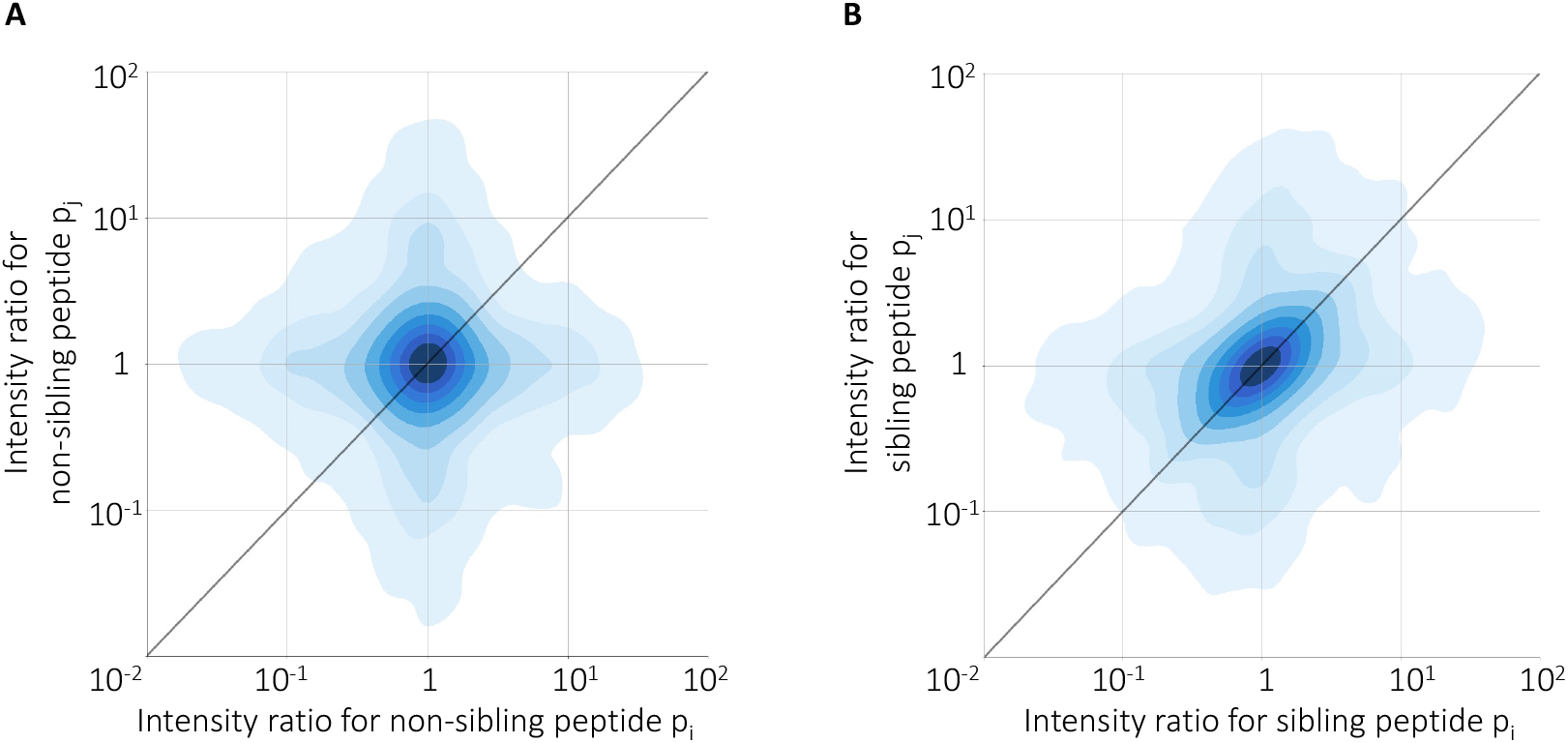
Consistency of sibling peptide ratios across experiments. The figure plots the log ratio of intensities of pairs of peptides across pairs of MS runs. Each point corresponds to a randomly selected pair of (A) non-sibling or (B) sibling peptides observed in a randomly selected pair of runs. Each panel contains 10,000 randomly selected pairs.

### 3.2 The model successfully generalizes to new peptides in new runs

Directly computing ratios of sibling peptide abundances is not a suitable strategy for inferring sequence-induced bias because the empirical ratios potentially reflect additional bias and noise. Accordingly, we turned to our neural network model, which learns to predict the peptide coefficient directly from the peptide sequence and charge state. We hypothesized that, if a Pepper model is truly learning sequence-specific biases, then the model should be able to generalize to new runs and new peptides. Accordingly, we segregated our data into a collection of training and test runs, and we similarly segregated proteins into train and test sets. We then trained a model using only training proteins drawn from the training runs, and we used the trained model to adjust the quantities associated with test proteins in the test runs.

The results of this experiment show that the model successfully generalizes (Figure 3). In particular, we find that the model reduces the loss—i.e., it succeeds in pushing the sibling peptide abundance differences closer to zero—in the test set. The kurtosis of the distribution of sibling peptide abundance differences is also reduced to from 6.01 to 5.18, highlighting the peak at 0 after adjusting with Pepper coefficients.

**Figure 3:**
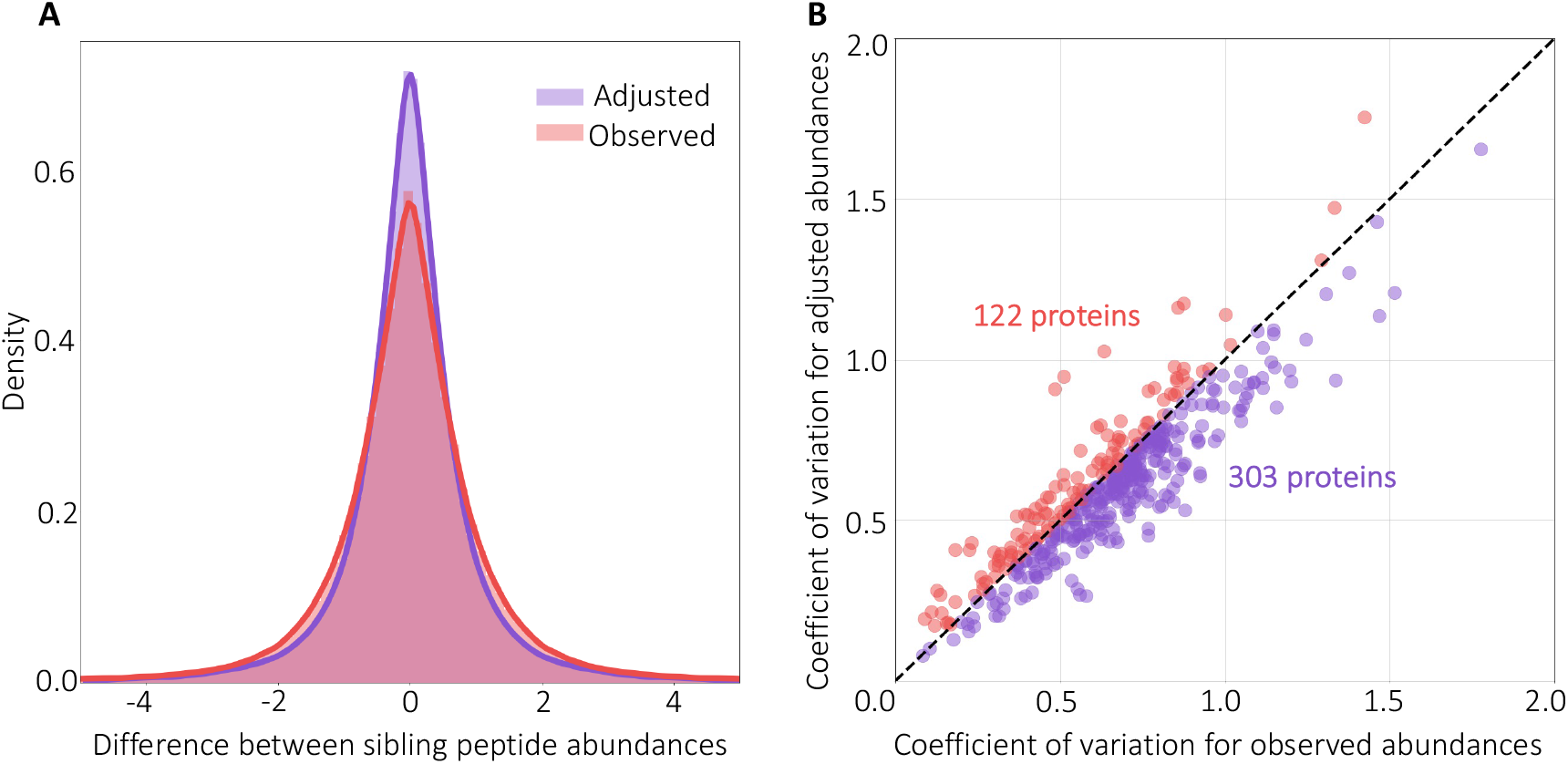
Predicting peptide coefficients across proteins and runs.

(A) The figure plots a histogram of the difference between the observed and adjusted peptide quantities of all sibling peptide pairs, on a logarithmic axis, for the test peptides and runs. (B) The figure plots the coefficient of variation (CV) before and after adjustment, for the test proteins and runs.

We repeated the training of the model 10 times with different random initializations and obtained an average loss reduction (i.e., Equation 1) of 28.71% in the test set. We also computed the coefficient of variation (CV) per protein (i.e., the standard deviation divided by the mean), before and after adjusting with Pepper coefficients, and observed that CV decreases for the majority (71.3%) of the test proteins (Figure 3B), highlighting the ability of the model to reduce the within-protein variance in test set abundances. To further examine the proteins which lead to an increase in the CV, we recorded the CV change in proteins with >5 peptides and observed that an even higher percentage of proteins had a decreased CV (80.8%).

To further visualize this result, we selected the 10 proteins from the test set with the highest number of quantified peptides and plotted the abundance of all peptides occurring on each protein before and after adjusting with the coefficients (Figure 4). In each case, we observe that the adjusted peptide abundances fall closer to the mean, indicating that our coefficients can minimize the bias in sibling peptide measurements for an unseen protein. We also report the per-protein change in loss values, alongside the change in CV for each protein. For all but one of the proteins, an improvement in the loss function corresponds to a decrease in CV. The only exception is protein O75533, where the increasing CV corresponds to one of the lowest loss changes. We further examined the results for this protein and found that a peptide with charge +6 is responsible for the high standard deviation; after excluding this peptide, the CV decreased by 25.4%. This observation matches with our expectation that the Pepper predictions are less reliable for high charge peptides, because our training set has only a small number of examples for higher charge states. These results support our two central hypotheses—that measured peptide quantities can be decomposed into a sequence-specific bias term and that the adjusted peptide quantities for sibling peptides should be approximately equal—and suggest that Pepper is able to learn to predict the sequence-induced bias terms. Note that, a priori, we do not expect the model to be capable of reducing the loss to zero, even on the training set, because the observed data presumably contains many biases that are not predictable from the amino acid sequence alone.

**Figure 4:**
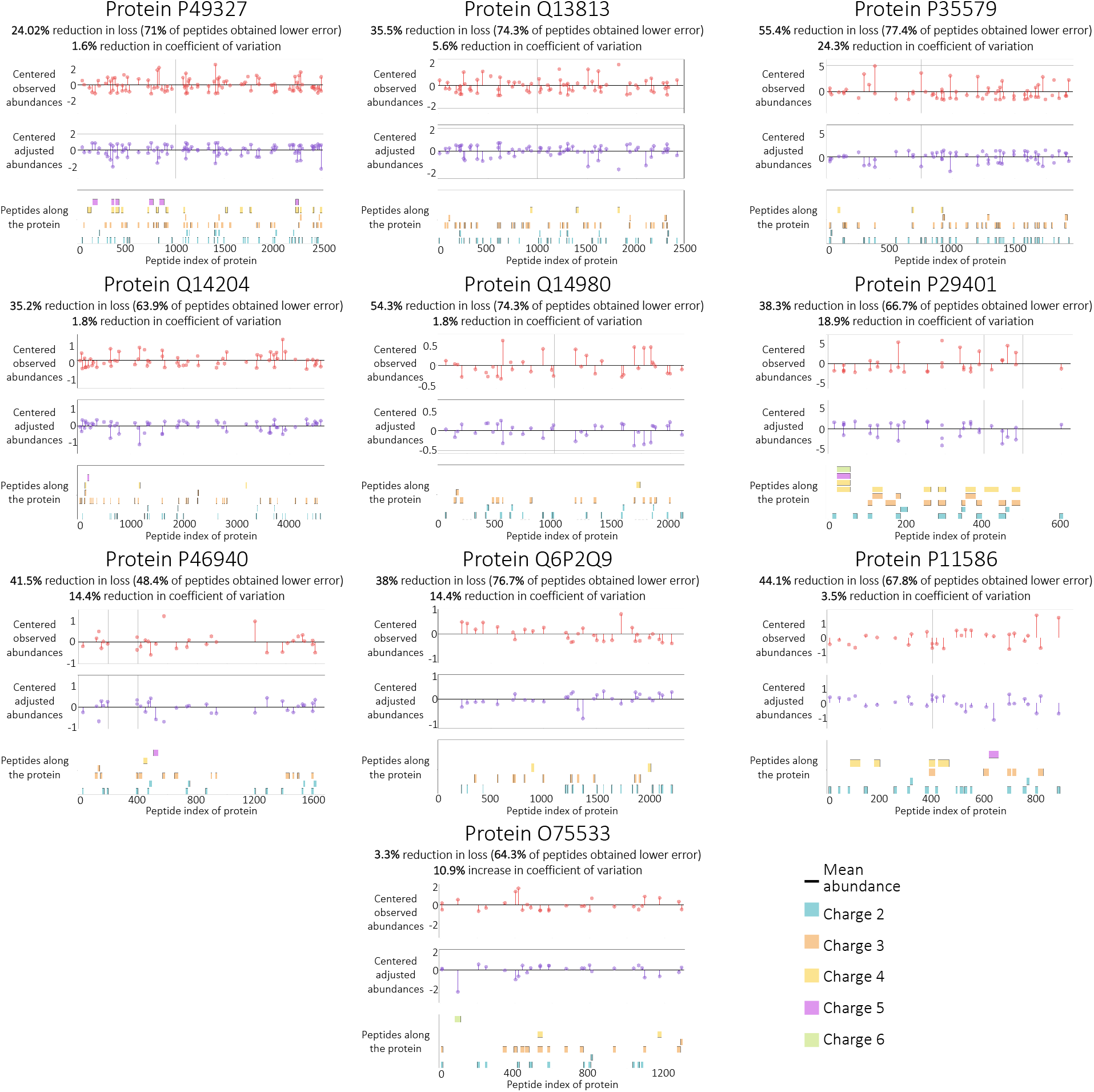
Comparing observed and adjusted abundances. The figure plots the abundance for all peptides occurring on 10 test set proteins with the highest numbers of peptides. The horizontal axis is amino acid position along the sequence, and the vertical axis shows the mean-centered peptide abundance for the original (top) and adjusted (bottom) abundances. The percent improvement is calculated with respect to the loss function (Equation 1). The percent reduction in coefficient of variation (CV) is also reported for each protein. Individual peptide sequences, segregated by charge state, are arrayed along the bottom of each figure.

We next tested whether this approach generalizes to other instrument types, acquisition strategies, and quantification schemes by training and testing the model using a variety of datasets (Table 3). In each case, we segregated the runs and proteins into train and tests sets in a ratio of 80% to 20%, and we used a fixed model architecture for training and testing. We observed that our model can successfully predict coefficients for datasets with different characteristics, in each case substantially reducing the test set loss relative to the baseline.

**Table 3:** Performance on various datasets. The table lists a variety of datasets, reporting in the final column the mean percentage reduction in loss on the test set, relative to the baseline, along with the standard deviation.

| Dataset | Instrument | Acquisition | Quantification | Improvement |
| --- | --- | --- | --- | --- |
| Slevlek *et al.* | 5600 TripleTOF | DIA | SWATH-MS | 33.00 ($\pm 4.31$) |
| Thomas *et al.* | 5600+ TripleTOF | DIA | SWATH-MS | 27.90 ($\pm 4.89$) |
| Guo *et al.* | 5600 TripleTOF | DIA | SWATH-MS | 28.71 ($\pm 4.10$) |
| CPTAC S16 | LTQ Orbitrap Velos | SIM | Label Free | 22.05 ($\pm 2.56$) |
| CPTAC S19 | LTQ Orbitrap Velos | DDA | Label Free | 9.55 ($\pm 5.05$) |
| CPTAC S37 | Q-Exactive Plus | SRM | Label Free | 15.83 ($\pm 0.24$) |
| CPTAC S47 | Orbitrap Fusion Lumos | DDA | TMT11 | 20.49 ($\pm 1.68$) |
| CPTAC S48 | Orbitrap Fusion Lumos | DDA | TMT11 | 14.88 ($\pm 1.91$) |
| CPTAC S49 | Q-Exactive HF | DDA | TMT11 | 17.50 ($\pm 4.57$) |
| CPTAC S51 | Orbitrap Fusion Lumos | DDA | TMT11 | 24.84 ($\pm 1.76$) |
| CPTAC S54 | Orbitrap Fusion Lumos | DDA | TMT11 | 15.31 ($\pm 0.65$) |
| CPTAC S58 | Orbitrap Fusion Lumos | DDA | TMT11 | 20.39 ($\pm 0.97$) |
| CPTAC S61 | Orbitrap Fusion Lumos | DDA | TMT11 | 20.30 ($\pm 1.79$) |

### 3.3 Pepper learns successfully in the presence of mislabeled proteoforms

One potential challenge faced by our model arises from the necessarily incomplete and inaccurate collection of proteoforms in our database. Although we train Pepper using a reference proteome that contains many known isoforms, the database cannot possibly account for the huge number of proteoforms that exist in a complex mixture, including unexpected isoforms, post-translational amino acid modifications, and truncation events. In practice, the incompleteness of our database will most often give rise to false positive labels in our training set, i.e., pairs of peptides that we believe to be siblings but which actually lie on different proteoforms (Figure 5A). To the extent that such false positives occur in real data, our model will be harder to train.

**Figure 5:**
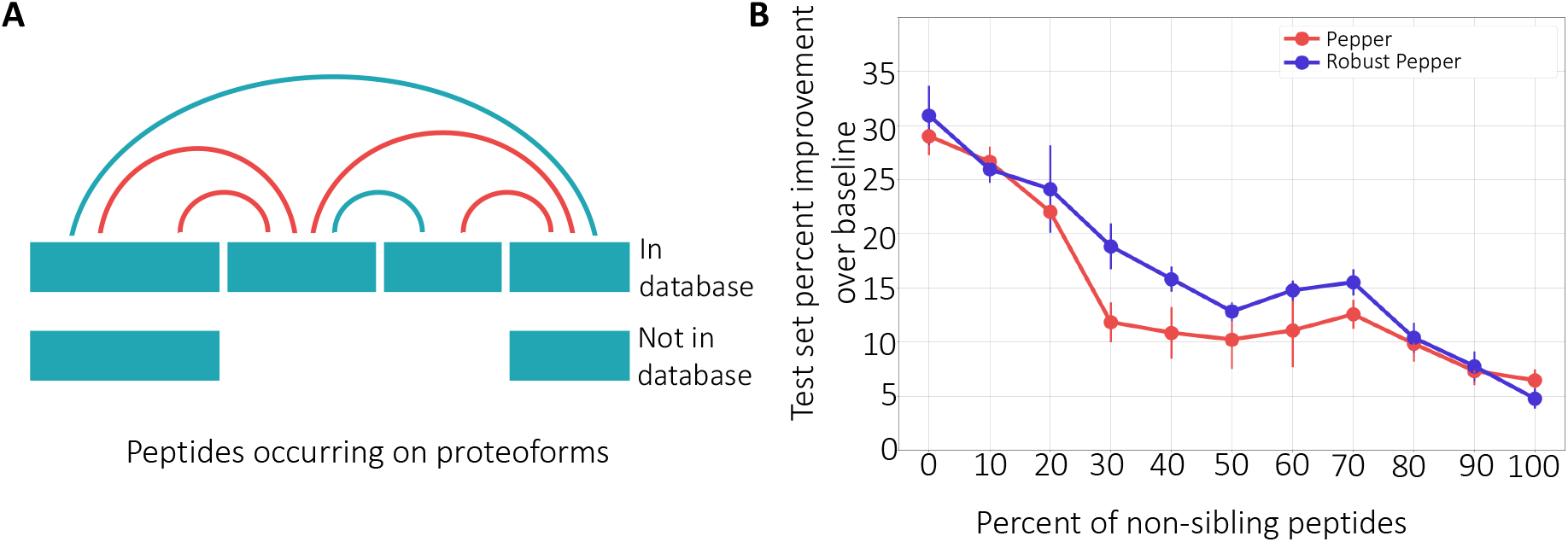
The model’s robustness to proteoform noise. (A) A protein that has two isoforms, only one of which is in the database. The four peptides yield six pairs of apparent siblings. However, because of the presence of the unknown isoforms, some of the sibling relationships (marked in red) are invalid. (B) The figure plots the percent improvement over the baseline (when all coefficients are set to 1), over ten runs, of a fixed set of test peptides as the percentage of label noise increases. Error bars correspond to standard error.

To investigate the robustness of our approach to proteoform noise, we artificially injected false positives into our training procedure and examined the behavior of the trained model. Specifically, we created a training set consisting of 14,472 peptides, and then we randomly permuted the protein labels for a fixed percentage of the training peptides. This procedure has the effect of creating false positive pairs, mimicking pairs of peptides that occur on a single proteoform in our database but occur on distinct proteoforms in the sample.

This experiment shows that the Pepper model is robust to such noise. We observe a smooth degradation of performance as the percentage of false positive pairs increases, with the improvement on the test set remaining above 10% even up to 70% false positives (Figure 5B). This result shows that even if our training set contained label noise, we are able to successfully learn to identify sequence-induced bias.

While we demonstrate the generalizability of our model in the presence of mislabeled sibling peptides, the test set performance is highly dependent on the number of noisy samples in the training set. Thus, we aimed to adapt our model to better handle label noise. Borrowing from a popular machine learning technique, Robust PCA [12], we extended our coefficient predictor to model the corrupted labels as parameters to be inferred. Specifically, we trained the same network using a modified loss function that is more robust to label noise:

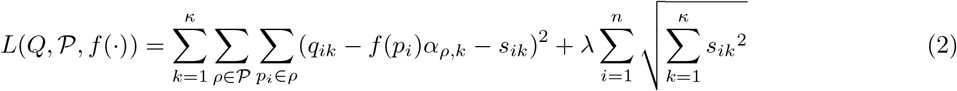

where *s_i,k_* is the noise term associated with peptide *i* and run *k*. The model is trained subject to the same constraints as the loss function in Equation 1. The second term in Equation 2 corresponds to the regularizer for the *S* matrix. The lambda value determines the strength of regularization, which we tuned using a validation set for each different noise ratio value.

We hypothesized that, by properly accounting for the label noise that we know exists in our data, this method will improve our ability to accurately infer peptide coefficients. Accordingly, we trained the robust model and compared the percent improvement across all noise ratios (Figure 5B). We observe that the robust model outperforms the regular model, indicating that the robust model is useful for eliminating label noise. We expect this extension of the model to be especially useful for datasets with high ratios of proteoform noise. On the other hand, this extension to the model significantly increases the number of parameters (by roughly a factor of 5 on average), making the training procedure significantly slower.

### 3.4 The coefficients reflect physicochemical properties of the peptide sequences

Previous machine learning models that aim to characterize peptide-specific biases in the context of identifying, rather than quantifying, peptides from mass spectrometry data have shown that the predictions from the models correlate strongly with several key peptide features, including hydrophobicity and peptide length [1–7]. Accordingly, we segregated Pepper’s coefficient predictions from our model according to these two features. In both cases, we observe a strong trend (Figure 6A–B), with coefficients taking smaller values for longer peptides or peptides with extreme values (high or low) of hydrophobicity, in agreement with previous work. These results indicate that the instruments are yielding under-estimates of the quantities of long or highly hydrophobic/hydrophylic peptides. By learning small coefficients for such peptides, our model aims to correct the associated biases.

**Figure 6:**
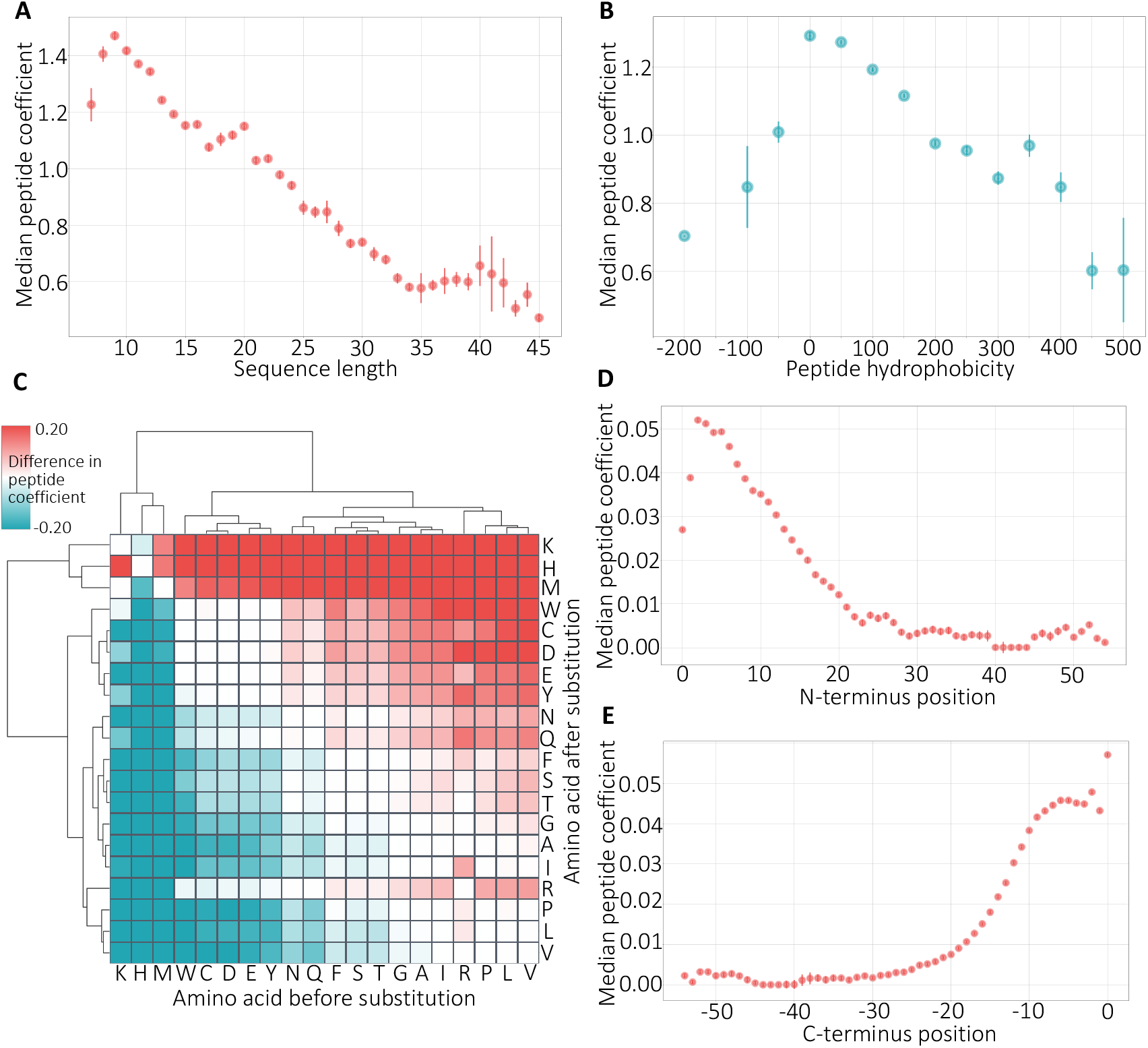
Physicochemical properties of peptides. (A) The figure plots the relationship between median peptide coefficient (y-axis) and sequence length. Bars represent standard error. (B) Same as panel A but for peptide hydrophobicity. (C) Cluster map of the change in the peptide coefficient per amino acid substitution. The values are the median over all instances of the given substitution in our simulation. (D) The figure plots the distribution of the change in the peptide coefficient plotted separately for each N-terminus position in the peptide sequence. (E) Same as panel D but for C-terminus positions.

We further calculated the correlation between the learned coefficients and 494 different physicochemical properties obtained from the AAIndex database [13]. Table 4 lists the highly correlated features, including polarity, hydration, and structural features. Polarity and hydrophobicity of a peptide determine its behavior in the solvent [2]. Structural features are also highly relevant because the structure can affect tryptic digestion [1]. It is promising to see our model capturing these properties of the peptides affecting how they behave in the mass spectrometer and adjusting their abundances to provide more accurate quantification.

**Table 4:** Top physicochemical peptide features. The table lists the peptide features yielding the highest correlation with the learned coefficients. All the features are obtained from the AAindex database [13]. For each feature, the table reports the absolute value of the Pearson correlation coefficient (|*r*|).

| Peptide feature | $\vert r\vert $ |
| --- | --- |
| Hydration number | 0.451 |
| Relative preference value at N3 (ends of alpha helices) | 0.442 |
| Relative preference value at N2 (ends of alpha helices) | 0.438 |
| Percentage of exposed residues | 0.437 |
| Side chain oriental preference | 0.437 |
| Alpha-helix indices for beta-proteins | 0.436 |
| Average accessible surface area | 0.436 |
| Polar requirement | 0.435 |
| Hydrophilicity scale | 0.434 |
| Helix initiation parameter | 0.431 |

While the previous methods relied on amino acid features summarized at the peptide level, they highlighted that amino acid composition is a potentially important feature affecting tryptic digestion [1, 2, 5]. Accordingly, we wanted to investigate the effect of amino acid substitutions on the peptide bias. To do so, we randomly sampled real peptide sequences from the NCI-60 dataset and generated simulated peptide sequences by substituting each amino acid at every position with every other amino acid. We then used our trained model to predict coefficients for all pairs of sequences and calculated the differences between the predicted coefficients.

The resulting clustered heat map (Figure 6C) suggests that the learned clusters align with physicochemical features of the amino acids. We observe that the two prominent clusters consist of mostly polar versus nonpolar amino acids. Polarity was among the most discriminative features for some previous approaches [1, 3, 5]. The polar cluster particularly consists of charged amino acids (D, E, H, K). The hydrophilicity and the charge of the residues affect fragmentation, ionization, and detection processes and thus are critical for determining the behavior of a peptide in mass spectrometer [1]. Similarly, the non-polar cluster contains a group of small amino acids (S, T, G, P, A) where the size of the side chain is known to be related to the flexibility of the amino acid [2]. The cluster map also highlights that the learned coefficients become larger, in general, when a hydrophobic amino acid is substituted with a polar one, indicating that more polar peptides are favored by the instrument. This finding agrees with studies that detected a negative correlation between hydrophobicity of the peptide and the probability of detection [6].

We also investigated the effect of the substitution position on the peptide coefficient. We grouped the scores by position with respect to the N- and C-terminus and plotted the distribution of the coefficients (Figure 6D–E). We find that amino acids at either end of the sequence are strongly related to sequence-induced bias. This might be because these residues play an important role in susceptibility of the peptide to enzymatic cleavage or absorption to solid phase extraction matrices [4]. Recapitulating the features that were shown to be important for determining the detectability of a peptide in the context of quantification highlights the biological relevance of the learned coefficients.

### 3.5 Pepper outperforms a simple linear model

The observed correlation between our predicted coefficients and amino acid composition suggests that perhaps a simple linear regressor trained using compositional features might be sufficient to accurately model peptide bias. To test this hypothesis, we trained a linear regression model from amino acid counts vector (i.e., vector of length 20 containing the number of occurences of each amino acid) trained using the same loss function as our neural network (Equation 1). We further trained linear regression models trained using 2-mer or 3-mer counts (i.e., vectors of length 400 or 8000 containing the number of occurrences of each *k*-mer).

The comparison of the models (Table 5) shows that Pepper clearly outperforms the alternative approaches, highlighting that the neural network architecture, which allows for nonlinearities and for dependencies between input features, is essential for the accurate prediction of the coefficients.

**Table 5:** Performance for baseline approaches. The table lists baseline approaches, reporting in the final column the mean percentage reduction in loss on the test set, relative to the baseline, along with the standard deviation.

| Model | Test set percent improvement |
| --- | --- |
| Pepper | 28.72 ($\pm 4.10$) |
| Amino acid (1-mer) count linear predictor | 13.30 ($\pm 5.46$) |
| 2-mer count linear predictor | 8.48 ($\pm 3.05$) |
| 3-mer count linear predictor | 15.03 ($\pm 1.24$) |

### 3.6 Factoring out sequence-specific bias improves correlation with gene expression

One of the criteria for evaluating the success of our approach is whether the adjusted abundances can provide more accurate quantification. We hypothesized that improving the accuracy of protein quantification would lead to higher correlation with the mRNA measurements. As has been discussed extensively in the literature, we do not expect a very strong correlation between these two data modalities, due to effects such as post-translation modifications and variations in protein degradation rates [14, 15]. Nonetheless, we reasoned that a small proportion of the discordance between protein and mRNA expression might be explained by sequence-specific biases in the quantitative proteomics data. Accordingly, we used a paired set of RNA-seq and mass spectrometry measurements to calculate the correlation between the gene and protein-level abundances for each sample before and after adjustment using Pepper. As expected, we observe that Pepper increases the correlation between protein and mRNA-based measurements (Figure 7A). Strikingly, the correlation improves in 59 out of 59 runs that we tested on (p < 1.2 × 10^−11^, signed-rank test) highlighting the ability of the learned coefficients to improve the accuracy of downstream analysis.

**Figure 7:**
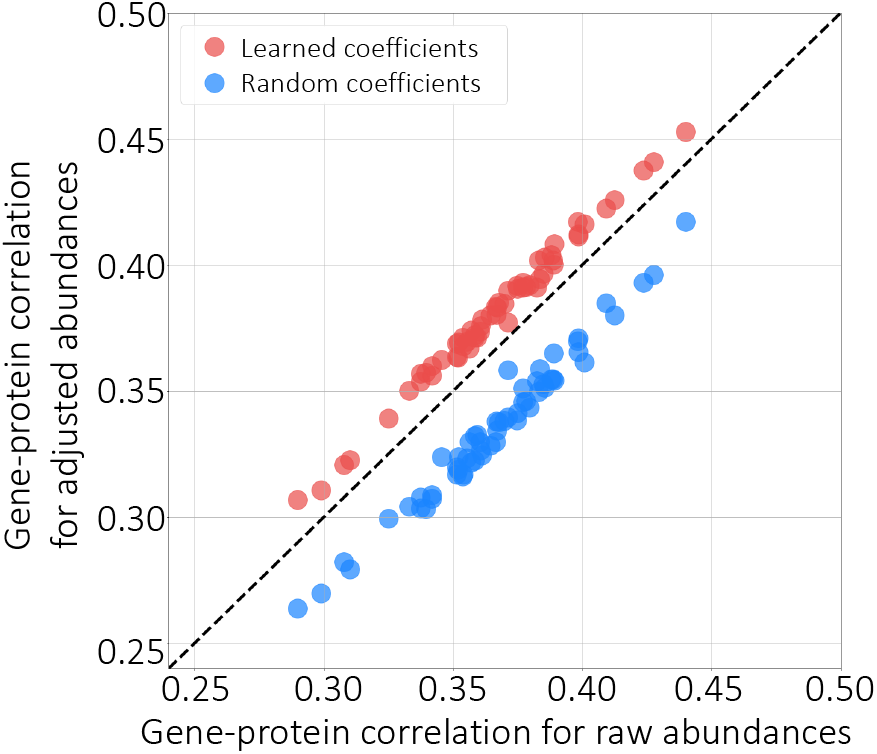
Factoring out sequence-specific biases. The figure plots the per-protein correlation between gene expression and protein expression, before (x-axis) and after (y-axis) adjusting the quantities using the deep neural network (red points) or using randomly selected coefficients (blue).

As a control, we further generated a random set of coefficients by sampling from the range of the learned coefficients. Adjusting the abundances with these random coefficients and recalculating the correlations with the mRNA measurements resulted in deterioration of correlation, highlighting the significance of our results (Figure 7). This analysis suggests that our coefficients can be used to reduce the biases associated with peptide measurements.

### 3.7 The model learns successfully from a few runs

Finally, we investigated the effect of the number of training runs on the model performance, and whether it is possible to reduce peptide bias using a few runs. Accordingly, we downsampled the training runs in our data and repeated the model training while recording the percent reduction on a fixed set of test runs. We observe the test set performance increasing as the number of training runs increase, as expected, indicating that training from a higher number of runs can be helpful in increasing the generalizability of the model (Figure 8). On the other hand, the Pepper model trained from only two runs can achieve a test set loss reduction of 27%, indicating that our coefficient predictor can learn effectively from a small number of runs.

**Figure 8:**
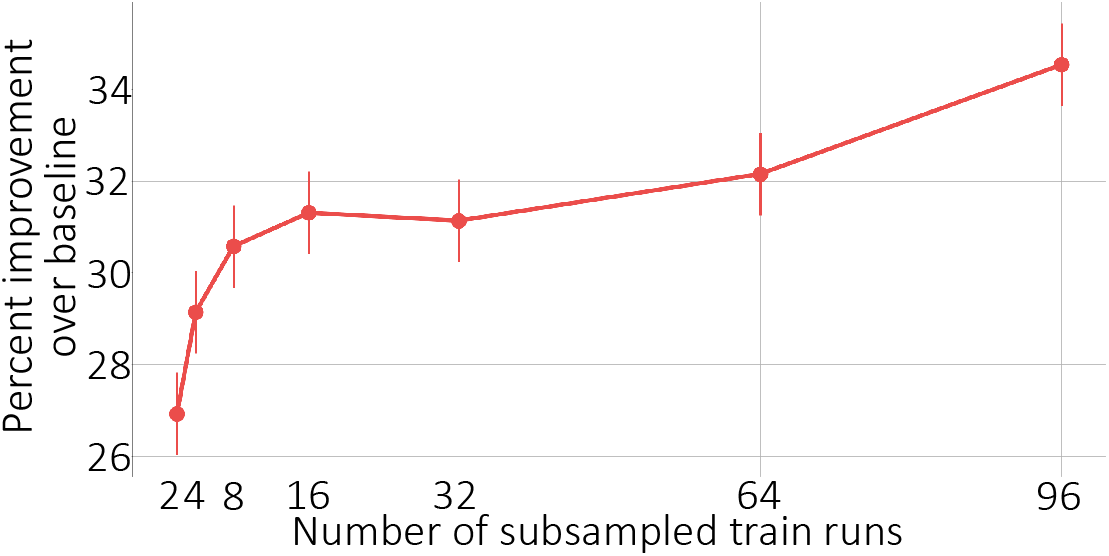
Peptide coefficient predictor learning curve. The figure plots the percent improvement over the baseline (when all coefficients are set to 1), over ten runs, of a fixed set of test peptides as the number of training runs increase, where 96 runs is the entire training set. Error bars correspond to standard error.

We also investigated Pepper’s ability to generalize across different mass spectrometry experiments. Accordingly, we trained our coefficient predictor on a combined set of all five CPTAC TMT11 datasets. We then applied the trained model to the held-out CPTAC TMT11 dataset (CPTAC S51) containing 35 runs.

This analysis showed a marked decrease in the model’s performance in the cross-experiment setting. In particular, when generalizing to new runs within the held-out dataset, the model achieved a reduction in loss of 24.84% (±1.76). In contrast, when generalizing to the held-out experiment, the improvement was only 13.51% (±1.45). The limited ability of our model to generalize across experiments might be related to experiment-specific biases associated with each dataset, which restricts our ability to transfer the learned coefficients to unseen datasets, as also observed in previous studies [1, 2].

## 4 Discussion

In this work, we aim to quantify the peptide-specific biases that arise in a quantitative MS/MS experiment, with the goal of adjusting the observed abundances to reduce bias. We developed a deep learning model, Pepper, that takes the peptide sequence and charge state as input and predicts a peptide coefficient to account for peptide-specific biases. Pepper was trained based on our assumption that the abundances of unique sibling peptides should be equal. We demonstrated that the predicted peptide coefficients successfully reduce our pre-defined loss function for new peptides and runs, which corresponds to reducing the CV of peptide intensities associated with a given protein. This generalization performance was replicated on multiple datasets generated with different MS/MS instruments using different acquisition and quantification techniques. We also detected significant correlation between various physicochemical features of peptides and the learned coefficients, highlighting that our model captures features, such as hydrophobicity and secondary structure, which were previously shown to affect how a peptide behaves in a mass spectrometer [1, 4–7]. We demonstrated that our coefficients significantly improve the correlation between protein and mRNA expression, and we examined Pepper’s ability to learn from datasets of varying sizes and to generalize across MS/MS experiments.

One caveat of our approach is that Pepper learns a single peptide coefficient to be applied across all runs. However, each mass spectrometry run exhibits specific biases. With our current approach, our analysis showed that the model pretrained on a different dataset—even if much larger—and transferred to the target dataset does not outperform a model trained directly on the target dataset itself. Previous studies also highlighted the same drawback, where the ability to predict across datasets was quite poor [1, 2]. Some of these studies even found that the most discriminative features for predicting the detection probability of a peptide changed from experiment to experiment [1, 2, 4]. Although training Pepper separately to learn distinct coefficients per run might seem like a plausible extension, our training procedure requires learning a function to map a peptide sequence to a generalizable coefficient. Hence, it is not possible to train our model using a single run. Alternatively, eliminating the run bias along with the peptide bias might be possible by learning run-specific peptide coefficients, but such a training scheme would require labels other than those produced using sibling peptide relationships. Such labels might, for example, be drawn from metadata about the run, embedded in the mzML header. Improving our model to overcome the dependency between the quantitative biases and the experiment would enable jointly training from hundreds of datasets and making predictions for new experiments. If this approach is successful, our ultimate goal would be to offer our trained model as a general preprocessor for any quantitative mass spectrometry data to improve downstream analysis.

In addition, even within a single MS/MS experiment, Pepper is currently limited to capturing only sequence-related biases. However, many additional biases exist that our approach is not designed to address, such as protease cleavage rates and effects of chromatographic elution. Properties of the protein structure may also be relevant, since burial or surface exposure of any given peptide depends on that structure. Thus, one future direction is to generalize our model to take into account a wider variety of features.

The Pepper software enables researchers with quantitative peptide measurements to infer coefficients and improve protein expression analysis. Better detection and understanding of bias in mass spectrometry experiments can change how we carry out and interpret experiments, leading to a better understanding of the proteome.

